# A novel hyper-parameter can increase the prediction accuracy in a single-step genetic evaluation

**DOI:** 10.1101/2022.07.03.498620

**Authors:** Mehdi Neshat, Soohyun Lee, Md. Moksedul Momin, Buu Truong, Julius H. J. van der Werf, S. Hong Lee

## Abstract

The H-matrix best linear unbiased prediction (HBLUP) method has been widely used in livestock breeding programs. It can integrate all information, including pedigree, genotypes, and phenotypes on both genotyped and non-genotyped individuals into one single evaluation that can provide reliable predictions of breeding values. The existing HBLUP method (e.g., that implemented in BLUPf90 software) requires hyper-parameters that should be adequately optimised as otherwise the genomic prediction accuracy may decrease. In this study, we assess the performance of HBLUP using various hyper-parameters such as *blending, tuning* and *scale factor* in simulated as well as real data on Hanwoo cattle. In both simulated and cattle data, we show that blending is not necessary, indicating that the prediction accuracy decreases when using a blending hyper-parameter < 1. The tuning process (adjusting genomic relationships accounting for base allele frequencies) improves prediction accuracy in the simulated data, confirming previous studies, although the improvement is not statistically significant in the Hanwoo cattle data. We also demonstrate that a scale factor, *α*, which determines the relationship between allele frequency and per-allele effect size, can improve the HBLUP accuracy in both simulated and real data. Our findings suggest that an optimal scale factor should be considered to increase the prediction accuracy, in addition to blending and tuning processes, when using HBLUP.

**Author Summary:** Despite significant advancements in genotyping technologies, the capability to predict the phenotypes of complex traits is still limited. H-matrix best linear unbiased prediction (HBLUP) method has been used to tackle this limitation to demonstrate a promising prediction accuracy. However, the performance of HBLUP depends heavily on the optimisation of hyper-parameters (e.g. blending and tuning). In this study, we introduce a scale factor (*α*), as a new hyper-parameter in HBLUP, which accounts for the relationship between allele frequency and per-allele effect size. Using simulation and real data analysis, we investigate the impact of the hyper-parameters (blending, tuning, and scale factor) on the performance of HBLUP. In general, the blending process may not improve the prediction accuracy for simulation and cattle data although a marginally improved prediction accuracy is observed with a blending hyper-parameter = 0.86 for one of carcass traits in the cattle data. In contrast, the tuning process can increase the HBLUP accuracy particularly in simulated data. Furthermore, we observe that an optimal scale factor plays a significant role in improving the prediction accuracy in both simulated and real data, and the improvement is relatively large compared with blending and tuning processes. In this context, we propose considering the scale factor as a hyper-parameter to increase the predictive performance of HBLUP.

## Introduction

Genomic prediction can achieve a relatively accurate prediction of additive genetic values or future phenotypes at an early life stage and has been applied in a broad range of disciplines, including animal breeding [1] and human disease risk prediction [2–4].

Genomic prediction requires genotypic information for both discovery and target samples. Genome-wide single nucleotide polymorphisms (SNPs) are typically used to estimate the genomic relationship matrix (GRM) for the genotyped samples so that breeding values (in livestock) can be estimated for the target samples, given the phenotypic information of discovery samples [5,6]. In many cases, we may have individuals with useful phenotypic information that are not genotyped, but they may be linked with genotyped samples through a pedigree, i.e., missing genotype data. To address this problem, a single-step genomic best linear unbiased prediction (ssGBLUP) method was introduced, in which phenotypic information on both genotyped and non-genotyped individuals in the pedigree can be used simultaneously to maximise the prediction accuracy of genotyped target individuals [7–9].

SsGBLUP uses an H-matrix that is a harmonised matrix of a pedigree-based numerator relationship matrix (NRM) and a GRM; therefore, we will use the term H-matrix best linear unbiased prediction (HBLUP). The H-matrix allows us to use the information of non-genotyped individuals in genomic prediction using a data augmentation technique (see [7, 8] and [10]). HBLUP has been widely used in the genetic evaluation of livestock and has been employed in the national genetic evaluation program in many countries [11–19]. There are numerous studies reporting that HBLUP outperforms traditional GBLUP [20–23].

In HBLUP, there are several hyper-parameters that can determine its performance. First, blending is one of the hyper-parameters that can provide a weighted sum of genomic and numerator relationships, using an arbitrary weight typically ranging from 0.5 to 0.99 [13]. This process is essential because it ensures GRM being a positive definite matrix to avoid numerical problems in HBLUP [7, 24]. Second, tuning is another important hyper-parameter that can adjust GRM, accounting for the allele frequencies in the base population that are inferred from the information of NRM [7, 8, 25, 26]. Note that GRM is typically based on genotyped samples in the last few generations, whereas NRM includes the information of founders in the base population through the pedigree. Third, a scale factor is a novel hyper-parameter for HBLUP, to be introduced in this study, which can generate different kinds of GRMs, accounting for the relationship between allele frequency and per-allele effect size, i.e. per-allele effect sizes vary, depending on a function proportional to [*p* (1 – *p*)]*^α^*, where *p* is the allele frequency [27–30]. Negative *α* values indicate lager effect sizes for rare variants, and the choice of *α* may determine the HBLUP accuracy, i.e., an optimal *α* can increase the accuracy.

In this study, we investigate for the three hyper-parameters, blending, tuning and *α*, to assess how they affect HBLUP accuracy, using simulated and real data. There are several tuning methods [7, 13, 25, 26] among which we test two most frequently used approach, i.e. methods by Chen et al. (2011) [26] and Vitezica et al. (2011) [25], referred to as tune=1 and 2 in this study. For blending, we investigate a wide range of weighting factor (*θ*), to assess the performance of HBLUP. In the analyses, we use the direct AI algorithm [31, 32] that is robust to the numerical problem caused by non-positive definite GRM so that we can assess all kinds of weighting factors in blending, including *θ* = 1. We also assess HBLUP performance, varying the scale factor, ranging from *α* = −1.5 to 1.5, in the estimation of GRM. We consider the three hyper-parameters simultaneously to obtain optimal values for blending, tuning and *α*, using a grid search method [33]. Then, the performance of HBLUP with the optimal values is compared to performances with less optimal values.

## Material and Methods

### Simulated data

QMSim software [34] was used for simulation since it can efficiently generate a large-scale dataset including genotypic and pedigree information. We simulated three different scenarios that differed in terms of the effective population size, mating design, and family structure.

I. The historical population consists of 100 generations. For the initial 95 generations, the effective population size (*N_e_*) keeps fixed at 100 individuals, consisting of 50 females and 50 males. Two offspring are generated with random selection and random mating of parents. In the following five generations (95^th^-100^th^), the number of progenies is gradually increased to 1000. In the last generation of the historical population (the 100^th^ generation), we select randomly 50 males and 500 females as the founders, and each male is mated with ten females and each female produced two offspring (i.e., a half-sib design). The current population consists of five generations with 1000 offspring in each generation (101 – 105^th^ generations), which is used for the main analyses. The details of applied parameters in the simulation of genotypic and pedigree data are listed in Table 1. The steps to simulate the historical and current populations are illustrated in S1 Fig.
II. In the second simulation scenario, *N_e_* = 1000 is used (500 females and 500 males) with a historical population of 100 generations. The population size for each generation in the historical population with 100 generations is constant (N=1000). In the subsequent five generations (101^st^ – 105^th^), each male is mated with one female and each female produced two offspring (i.e., a full-sib design) and 1000 offspring were generated in total. Thus, the founder population size is 1000.
III. In the third scenario, *N_e_* and the number of generations in the historical population are the same as the first scenario (*N_e_*=100 with 100 generations). However, In the last generation of the historical population (100^th^) and the subsequent five generations (101^st^ – 105^th^), the mating design and family structure are the same as the second scenario, i.e. one male is mated with one female to produce two progeny per mating (full-sib design), producing 1000 offspring in total in each generation.

**Table 1.** Parameters of historical population and genotyping data simulation in the first scenario using QMSim software.

| QMSim parameters | value |
| --- | --- |
| Litter size | 2 |
| Proportion of male progeny | 0.5 |
| Mating design | random (*rnd*) |
| Selection design | random (*rnd*) |
| Number of SNPs | $9 \times 10^3$ |
| Number of Chromosome | 30 |
| Chromosome length (cM) | 100 |
| Marker positions | random (*rnd*) |
| Marker allele frequencies | *eql* |
| Marker mutation rate | $2.8 \times 10^{-8}$ |

In order to simulate the phenotypes of a complex trait, based on the simulated genotyped data, we used a model,

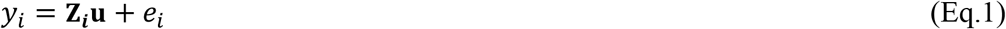

where *y_i_* is the phenotypic value, **Z_i_** is the vector of SNP genotypes and *e_i_* is the residual effect for the *i^th^* individual, and **u** is the vector of SNP effects. In this phenotypic simulation, we randomly selected 1000 SNPs as causal variants, and **u** was drawn from a normal distribution such that the mean and variance of the genetic effects are *mean*(**Z*_i_*u**) = **0** and *var*(**Z*_i_*u**) = *h^2^*. The residual effects were generated from a normal distribution with mean = 0 and variance = 1 – *h*^2^. In the phenotypic simulation, the SNP effects, **u,** are scaled by [2*p* (1 – *p*)]*^α^*, considering a non-negligible relationship between allele frequency and per-allele effect size [27–30], which is a function of alpha ranging from −1.5 to 1.5 in the simulation.

In the HBLUP analysis, for three simulation scenarios, it is assumed that the pedigree information is available for the last five generations (101 – 105^th^ generations), and the genotypic information is available for the individuals from the last two generations (104 – 105^th^ generations), noting that the sample size in each of the last 5 generations is 1000.

### Real data

#### Hanwoo cattle data

In this study, we applied statistical analyses to genotypic and phenotypic data from Hanwoo beef cattle. The total number of animals with pedigree information was 84,020, and among them, 13,800 animals were genotyped for 52,791 genome-wide SNPs, and 25,502 animals were recorded for their phenotypes. The number of animals available for both genotypic and phenotypic information was 9,072. The following criteria were applied for QC using PLINK: minor allele frequency below 0.01 (MAF), filtering SNPs with call rate lower than 95% (GENO = 0.05), individual missingness more than 5% (MIND= 0.05), and Hardy–Weinberg Equilibrium P-value threshold lower than 1e-04 (HWE). After QC, the number of individuals did not change, and SNPs number was 42,795. The Hanwoo beef cattle data included five carcass traits: carcass weight, eye muscle area, back fat thickness, marbling score and adjusted 12 months weight. The total number of animals with non-missing records for each carcass trait with and without genotypic information can be seen in Table 2.

**Table 2.**
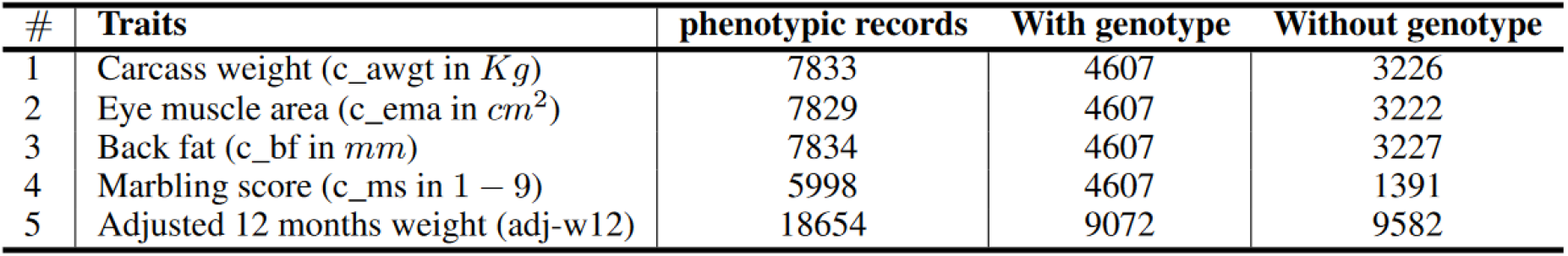
The number of individuals available for phenotypes with and without genotypic information for five carcass traits in Hanwoo cattle dataset.

| # | Traits | phenotypic records | With genotype | Without genotype |
| --- | --- | --- | --- | --- |
| 1 | Carcass weight (c_awgt in *Kg*) | 7833 | 4607 | 3226 |
| 2 | Eye muscle area (c_ema in *cm*<sup>2</sup>) | 7829 | 4607 | 3222 |
| 3 | Back fat (c_bf in *mm*) | 7834 | 4607 | 3227 |
| 4 | Marbling score (c_ms in 1 – 9) | 5998 | 4607 | 1391 |
| 5 | Adjusted 12 months weight (adj-w12) | 18654 | 9072 | 9582 |

In the HBLUP analysis for the Hanwoo cattle data, animals available for phenotypes and genotypes (*N_g,p_*) (see Table 2) are randomly divided into five groups. In a five-fold cross-validation, one of the five groups is selected as the target dataset, and the remaining groups are used as the discovery dataset, which is repeated for five times and the average prediction accuracy is achieved. The technical details of training and validating of HBLUP can be seen in Fig 1.

**Fig 1.**
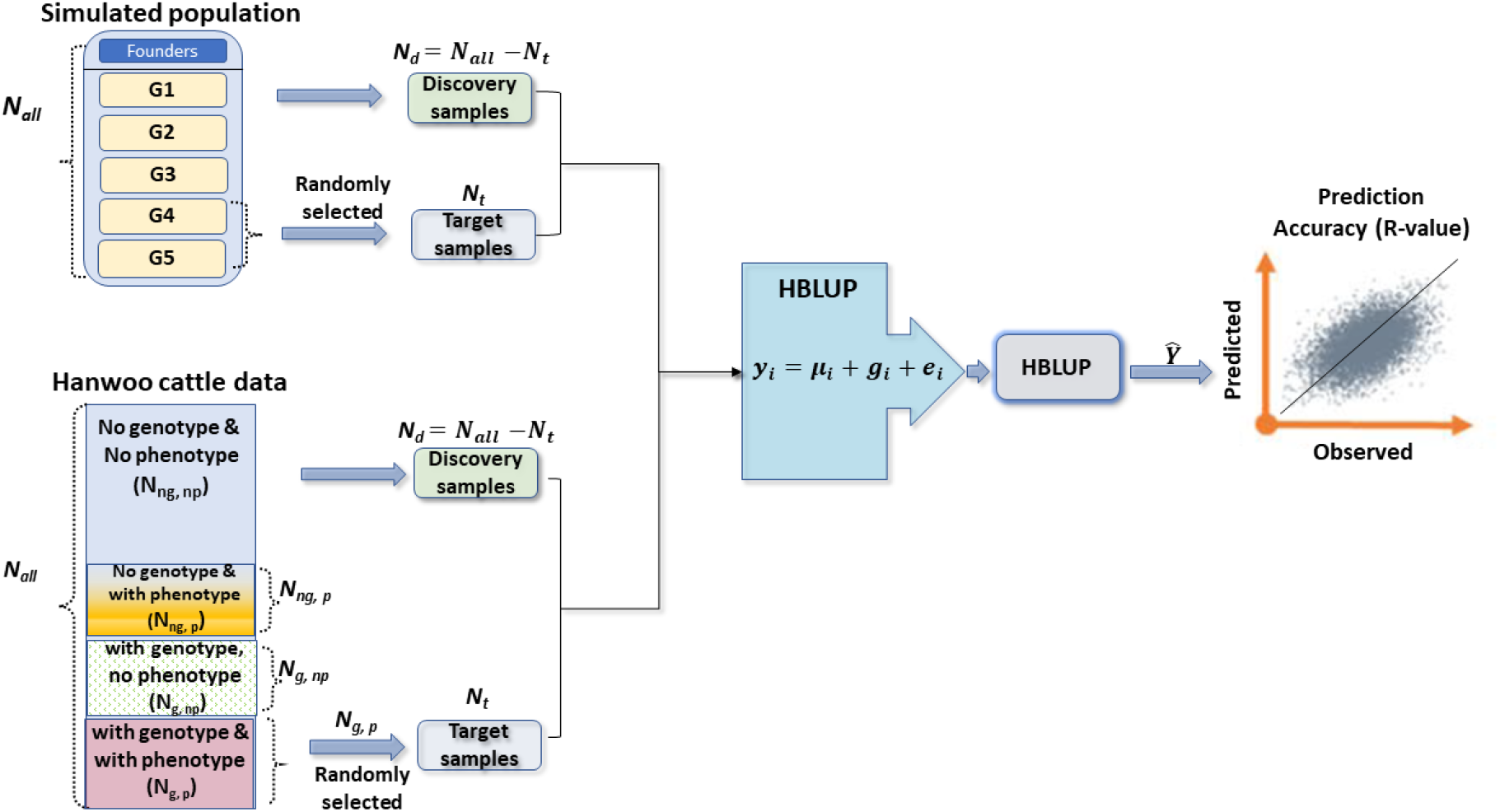
A diagram showing the experimental designs how to select the target and discovery samples for simulated and Hanwoo cattle datasets. In simulated dataset, the number of founders depends on the simulation scenarios (*f_n_* = 550, 1000 and 550 for simulation scenario 1, 2 and 3). The sample size in each generation (*G_i_*) is 1000. Therefore, the sample size in the whole population is 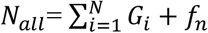. The sample sizes of target and discovery samples are denoted as *N_t_* and *N_d_*. In Hanwoo cattle data, the phenotypic and genotypic information is partly missing. The numbers of animals without genotype and phenotype (*N_ng,np_*), animals without genotype but with phenotype (*N_ng,p_*), animals with genotype but without phenotype (*N_g,np_*), and animals with both genotype and phenotype (*N_g,p_*) are shown in the diagram. *N_g_* is the total number of genotyped animals. In HBLUP, for the animals with both genotype and phenotype (*N_g,p_*), 5-fold cross validation is applied, and each fold is selected as the target dataset (*N_t_*), and the remaining animals with phenotypes are used as the discovery samples (*N_d_*). The best linear unbiased predictions for the phenotypes of the target samples are obtained. In order to calculate the prediction accuracy, we used Pearson’s correlation coefficients between the true and predicted phenotypes for the target samples.

#### Estimating NRM, GRM and HRM

##### Numerator relationship matrix

NRM denotes as **A** that is estimated based on the pedigree, which has been used in Henderson’s mixed model equation (1975) [35] to obtain estimated breeding values. Following [10], **A** matrix can be formulated as follows.

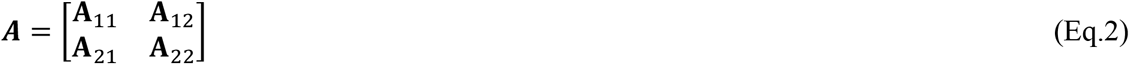

Where **A**_11_ and **A**_22_ denote the numerator relationships for the groups of non-genotyped and genotyped individuals, and **A**_12_ and **A**_21_ are the numerator relationships between non-genotyped and genotyped individuals.

#### Scale factor (*α*) and GRM

Following [29], the variance of the *i^th^* genetic variant (*v_i_*) can be expressed as a function of the allele substation effect (*u*) and the allele frequency (*p_i_*), which can be written as

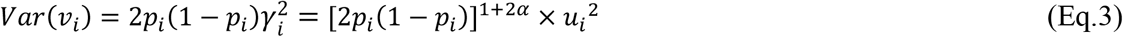

where *γ_i_* = *u_i_* × [2*p_i_* (1 – *p_i_*)]*^α^* is the allele effect size (**u***_i_*) that can vary, depending on the allele frequency and the scale factor, *α* [27, 28], which can be explained by evolutionary forces such as selections, mutations, immigrations, and genetic drift. In the classical model [36], *α* is assumed to be zero for all traits. Another widely used *α* value is *α* = −0.5, assuming that the genetic variance of the causal variant has a uniform distribution across the minor allele frequency spectrum. However, there have been reported that optimal *α* values vary, depending on traits and populations ([27, 28, and 29]).

Following [37], the genomic relationship matrix can be formulated as a function of *α*, which can be written as

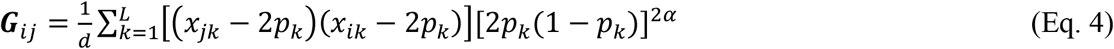

where ***G**_ij_* is the genomic relationship between the *i^th^* and *j^th^* individuals, and *L* is the total number of SNPs, *p_k_* is the allele frequency of the *k^th^* SNP, *x_jk_* is the SNP genotype coefficient of the *j^th^* individual at the *k^th^* SNP, and *d* is the expected diagonals computed as 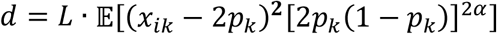. This Eq. 4 is implemented in LDAK software [27].

Note that Eq. 4 with *α* = −0.5 is equivalent to the genomic relationship estimation implemented in PLINK, GCTA and option 2 in BLUPf90 [24, 38, 39], and Eq. 4 with *α* = 0 is equivalent to option 1 in BLUPf90 [24, 38].

In the HBLUP analysis, we will vary *α* from −1.5 to 1.5, to find an optimal *α* value that can improve the prediction accuracy and compare the performance with the conventional HBLUP (with *α* = −0.5 or 0).

#### H-matrix best linear unbiased prediction

In the HBLUP analysis, GRM (**G**) is computed based on the genotypic information, and NRM (**A**) is estimated using the pedigree information of the population. Following [7], given estimated **G** and **A** (from Eq. 3 and 4), **H** matrix can be derived as

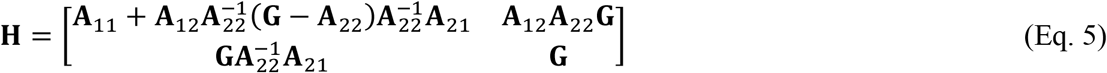

In the HBLUP analysis, the simulated data was divided into two groups, one group included the individuals in the first three generations, and the other group included individuals in the last two generations in the current population (101 – 105^th^ generations). We used the genotypic information of the last two generations and the full pedigree information across the five generations to estimate **H** matrix. In cattle data, animals available for phenotypes and genotypes were considered (see Table 2) to estimating GRM, and then the HRM was estimated using a combination of NRM estimated based on whole pedigree (84,020 individuals) and GRM.

#### Blending

GRM is typically a non-positive definite matrix. In the process of HBLUP, it is usually required to modify GRM to be positive definite so that it can be inverted without any numerical problem [24]. This modification method is called *‘blending’* that shrinks the genomic relationships toward the pedigree relationships, using an arbitrary weight, *θ*, typically ranging from 0.5 to 0.99 [13, 24]. The blended GRM can be written as

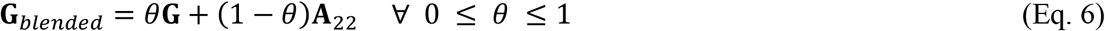

#### Tuning

Tuning process adjusts GRM, accounting for the allele frequencies in the base population, using the information from NRM that includes the information of founders in the base population through the pedigree [7, 8, 25, 26]. The tuned GRM (**G***_tuned_*) is computed as

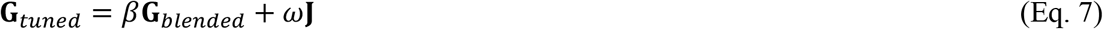

where **J** is a matrix with the same size of GRM, and all elements are equal to one, and *ω* and *β* are tuning parameters that can be used to adjust GRM, accounting for base allele frequencies. In this study, we use two most frequently used methods to obtain the tuning parameters, *ω* and *β*. Following [26], the first method (referred to as tune=1) computes *ω, and β* as

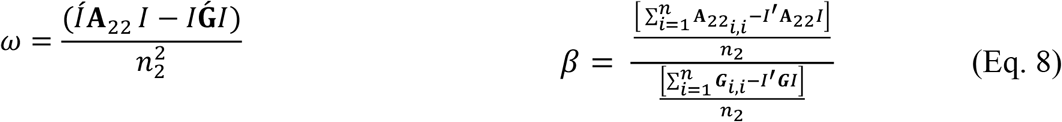

where **I** is an array with the size of *n* × 1 and all values equal to one.

Following [25], the second method (referred to as tune=2) can be written as

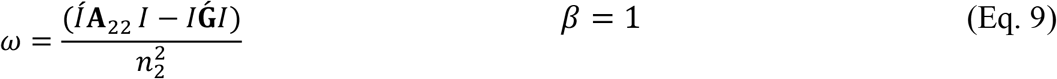

Please note that Eqs. 8 and 9 have been implemented in BLUPf90 [38] as the second and fourth tuning option (i.e. TunedG=2 or 4).

#### Linear Mixed Model

In the analyses, we used a linear mixed model that can be written as

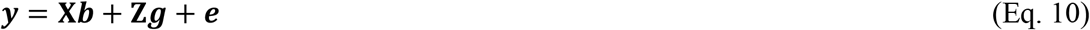

where ***y*** denotes a vector of phenotypic value, ***b*** is a vector of the (environmental) fixed effects, ***g*** is a vector of random additive genetic effect that is distributed based on 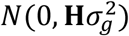, where **H** can be derived from Eq. 5 and 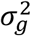 denotes the genetic variance. Both X and Z are the incidence matrixes. Finally, the residual effect vector is shown by ***e*** distributed as 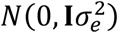 where **I** is an identity matrix and 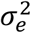 is the residual variance.

We employed the restricted maximum likelihood (REML) method, fitting the **H** matrix, to estimate genetic variance and heritability, which is referred to as HREML in this study. The Akaike Information Criterion (AIC) was used to assess the goodness of fitness of the model as *AIC* = 2*P* – 2 × ln(*L*), where ln(*L*) is the log likelihood from HREML, and *P* is the number of parameters. Given the estimated variances and heritability from HREML, HBLUP was used to obtain individual genetic values. We used MTG2.22 [44–45] genomic analysis software to perform HREML and HBLUP methods.

#### Grid Search to find optimal hyper-parameters

One of the well-known methods to find the best configuration of hyper-parameters is the grid search [40]. In the grid search, all possible combinations of hyper-parameters are considered to evaluate the performance of prediction models.

## Results

### Simulated data

Fig 2a shows that the tuning process significantly improves the prediction accuracy (referred to as R-value) that is a Pearson correlation coefficient between the observed and predicted phenotypes in the target dataset, confirming previous studies, when using the simulated data.

**Fig 2.**
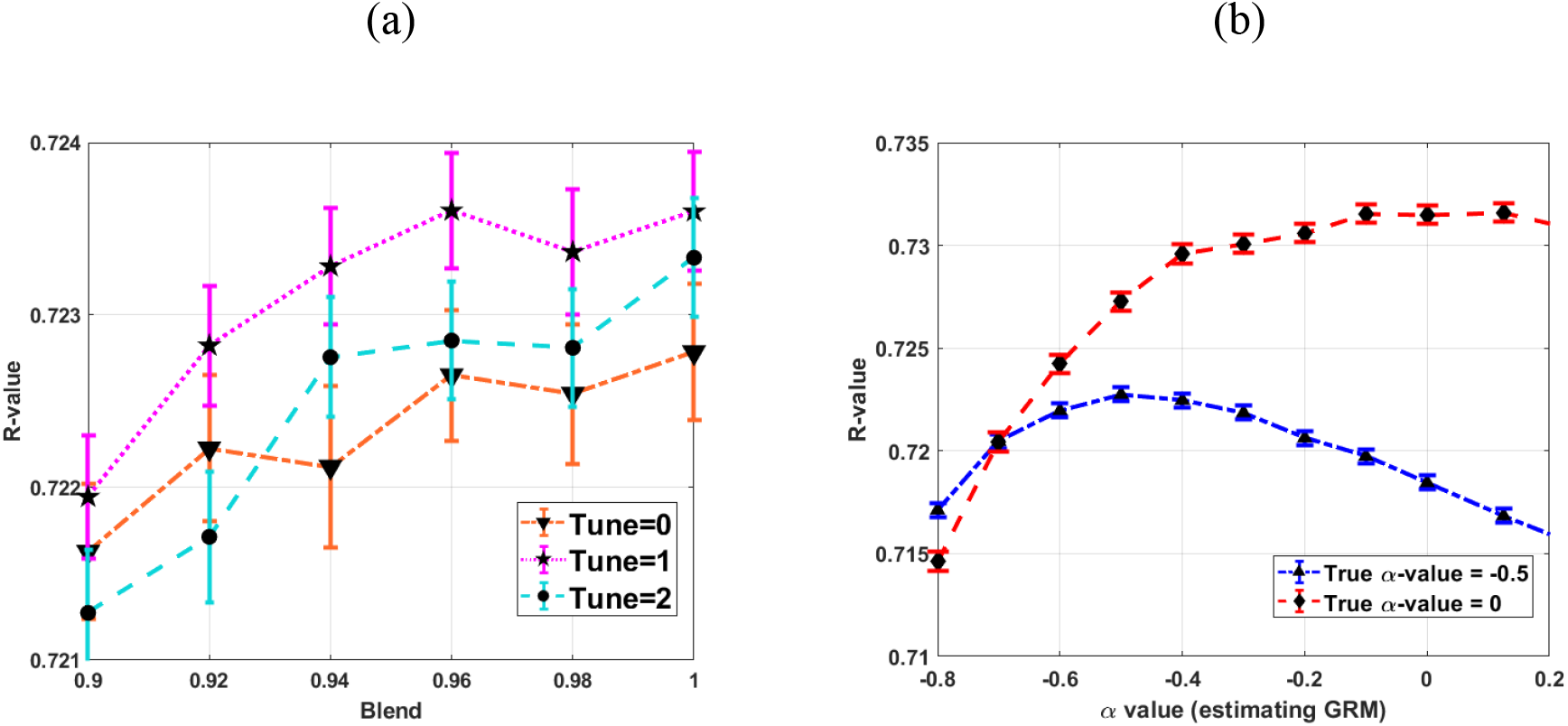
HBLUP accuracy and hyper-parameters. (a) The HBLUP accuracy (R-value) improves when using tune=1 (Eq. 8) or tune=2 (Eq. 9). However, blending (*θ* < 1) would not increase the accuracy for this simulated dataset. (b) Optimal *α* values can increase the accuracy, indicating that the choice of *α* is important in HBLUP. We simulated genotypes and phenotypes in 3000 replications in which simulation parameters of *h^2^* =0.8, *N_e_* = 100 for 100 historical generations and a half-sib design (50 male, 500 females) were used. The true *α* values used in the phenotypic simulation were −0.5 or 0. The error bars are 95% CI over the 3000 replications.

The tuning process with the first option (tune=1; Eq. 8) appears to better perform than the second option (tune=2; Eq. 9) for this simulated data. However, blending (*θ*<1) does not significantly improve the HBLUP accuracy for this simulated data (Fig 2a; S2 Fig). Fig 2b represents the impact of *α* value on the HBLUP’s performance, showing that the prediction accuracy increases when *α* value used in estimating GRM is close to the true *α* value used in the phenotypic simulation. When varying simulation scenarios (e.g., a small or large effective population size with full-sib designs), a similar result is observed that the prediction accuracy improves when applying the tunning process or when using optimal *α* (S3 Fig; S4 Fig; S5 Fig; S6 Fig).

Mimicking a real dataset in which multiple replicates are not possible, we used a single simulation data to assess the HBLUP accuracy, varying hyper-parameters (Fig 3). All possible configurations of tuning, blending and *α* values were evaluated using the grid search method where the prediction accuracy was measured using 5-fold cross validation (see Methods, S7 and S8 Figs). Fig 3 shows the HBLUP accuracy averaged over 5-fold cross validation when varying hyper-parameters. The highest prediction accuracy was achieved with tune=1, blend=1 and *α* =0 when using the true *α* =0, and with tune=1, blend=0.9 and *α* =-0.5 when using the true *α* =-0.5 in the simulations (See Fig 3). This shows that the optimal *α* values found in the grid search are approximately agreed with the true simulated values.

**Fig 3.**
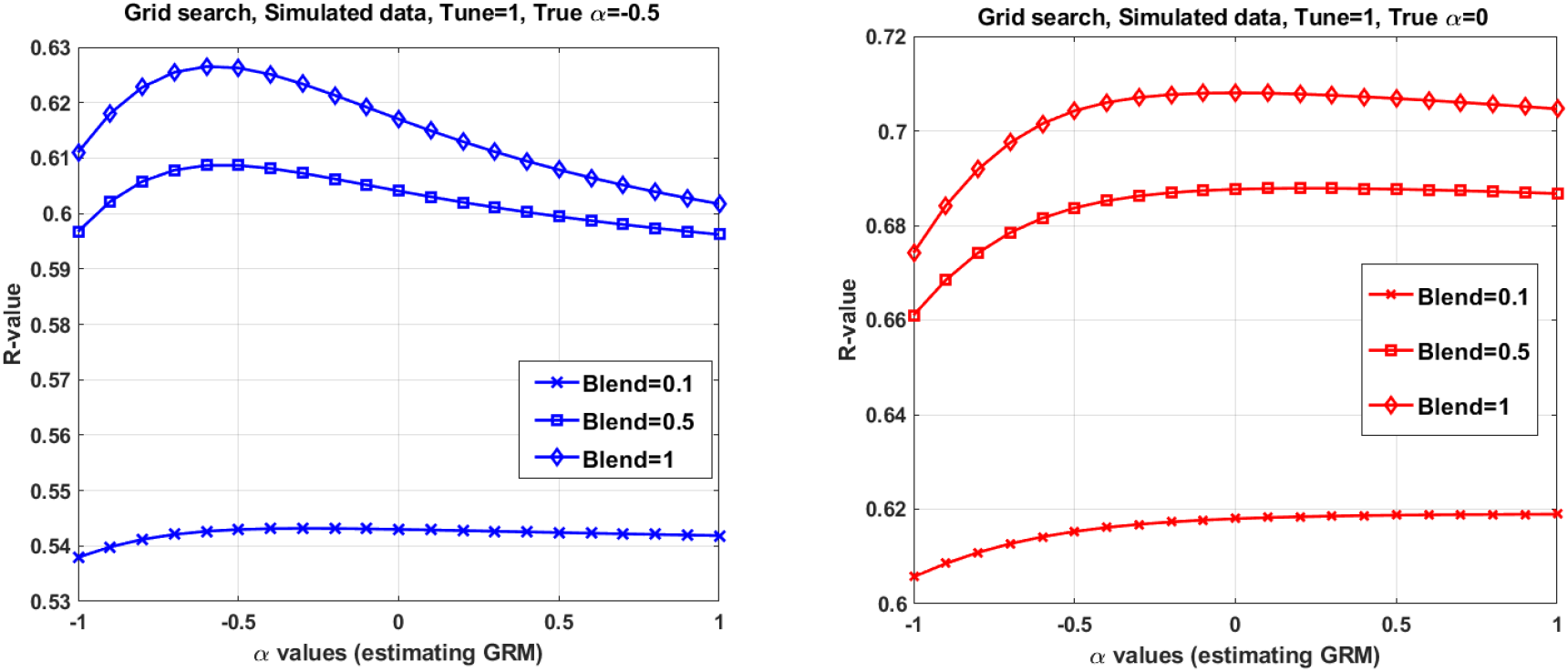
HBLUP accuracy averaged over 5-fold cross validation in a grid search with various configurations of the hyper-parameters, using a single simulation dataset. The best configuration found in the grid search consists of tune=1, blend=1 and *α* =0 (in estimating GRM) when using *α* =0 in the simulation, and tune=1, blend=0.9, and *α* =-0.5 when using *α* =-0.5 in the simulation. The population parameters used in the simulation are *h*^2^ =0.8, *N_e_* = 100 for 100 historical generations, NSNPS = 9000, chromosome number = 30 and *α* = 0 or −0.5. Mimicking livestock population, a half-sib design (50 sires, 10 dams per sire and 2 offspring per dam) was applied to the last 5 generations. Full pedigree across the 5 generations were used in HBLUP. Among 2000 offspring in the last 2 generations, 5 subsets each with a random 400 individuals were used as target datasets in the 5-fold cross validation. To predict for each target dataset, the remaining 5150 (across the 5 generations) were used as the discovery dataset.

#### Cattle data

We used pedigree, genotype and phenotype data of Korean native cattle (Hanwoo), which is a unique and important breed in the beef industry [42–43], to assess the HBLUP accuracy with various hyper-parameters including *α*. We first estimated optimal hyper-parameters that provided the lowest Akaike information criteria (AIC) value based on the residual maximum log-likelihood for each trait, using HREML (Fig 4). We observed that Δ*AIC* was not uniformly distributed across different *α* values, and optimal *α* values were largely different across 5 carcass traits (Fig 4a). On the other hand, a blending parameter *θ* = 1 provided the lowest Δ*AIC* values for all traits except of EMA (*θ* = 0.86), indicating that a blended GRM with *θ* < 1 did not increase the goodness of fit when using HREML in general (Fig 4b). Finally, Fig 4c shows that tune=2 could achieve a better goodness of fit, compared with tune=1 or tune=0 (i.e., without tuning), in most cases. For BFT and MS traits, tune=1 and 0 provided the lowest AIC (Fig 4c) although the AIC was not significantly lower than tune=2 (difference in AIC less than 1). The best-performed hyper-parameters for five traits can be seen in S1 Table.

**Fig 4.**
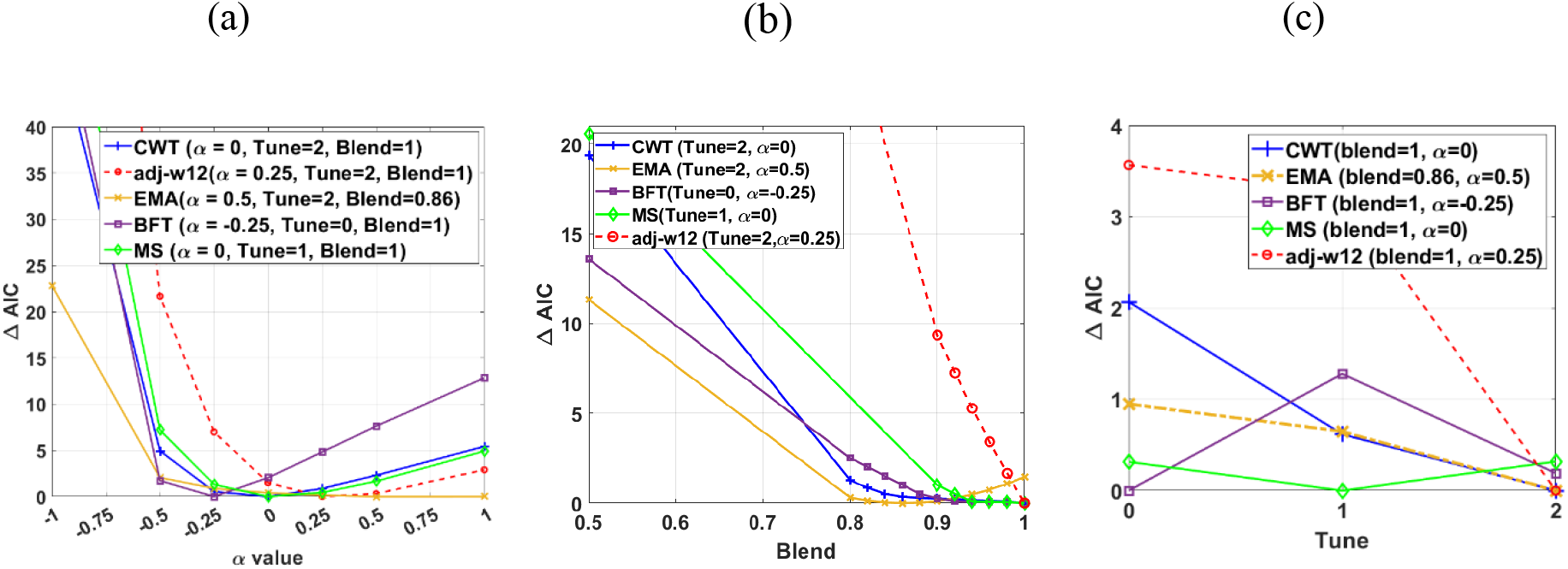
HREML estimation accuracy depending on *α* estimated in the genotyped samples and making HRM. (a) Evaluating the impact of *α* values on the Δ*AIC* for five different traits of Hanwoo cattle dataset using HREML in a univariate linear mixed model with different tuning methods and blending coefficients. The Akaike Information Criterion (AIC) was used to show the goodness of fitness of the model as *AIC* = 2*P* – 2 × ln(*L*), where 2 × ln(*L*) is the HREML log likelihood, and *P* is the number of parameters. Δ*AIC* = *AIC* – *AIC_optimal_*, where AIC is obtained with the corresponding *α* value at the x-axis and *AIC_optimal_* is the AIC for the optimal *α*. It is observed that optimal *α* varies across traits. Whole individuals with available phenotype were applied in estimating the heritability based on Table 2. (b) A performance comparison between two different blending coefficients (0.5 to 1) in order to estimate the HRM using HREML with optimal tuning method and optimal *α* value. (c) The performance of tune=1 (Eq. 8) compared with the tune=2 (Eq. 9) and without considering the tuning in estimating the HRM with the applied optimal blending and *α* values.

We also used a grid search to assess the performance of all hyper-parameters (Fig 5) in which HBLUP accuracies of all possible configurations of tuning, blending and *α*. values were evaluated in 5-fold cross validation. Fig 5 shows the HBLUP accuracy averaged over 5-fold cross validation when varying *α*, tuning and blending values for 5 carcass traits. In Fig 5a, we observed that the accuracy of HBLUP could be considerably increased or decreased, depending on the choice of *α* values. In contrast, Fig 5b shows that the highest HBLUP accuracy was achieved with a blending parameter *θ* = 1 for all traits except EMA (*θ* = 0.86), indicating that blended GRM would not improve the HBLUP accuracy in most cases. Finally, Fig 5c indicates that tuning process would not substantially improve the HBLUP accuracy for all carcass traits in Hanwoo cattle data. The best configuration of the hyper-parameters for each trait is shown in S1 Table.

**Fig 5.**
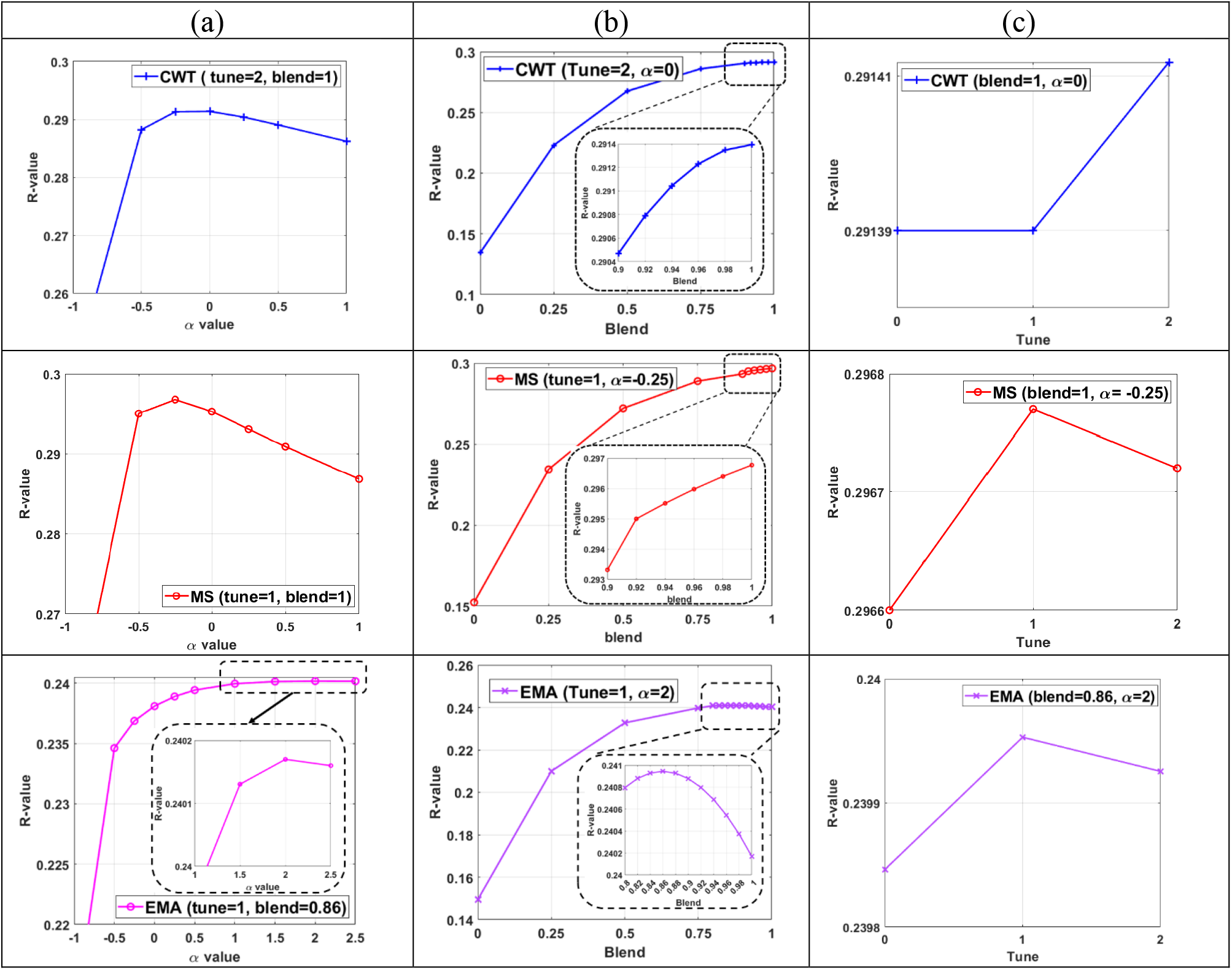
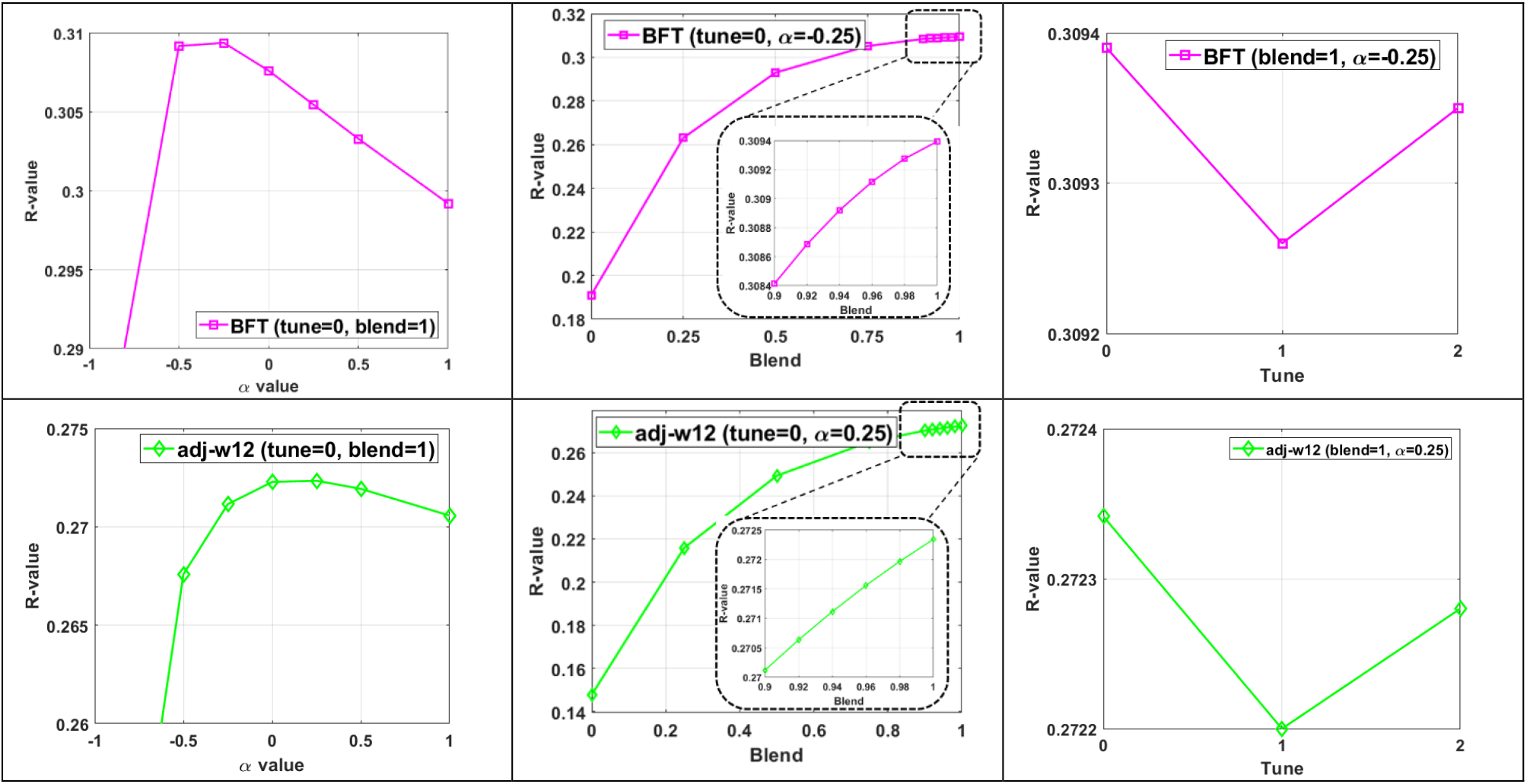
The performance of HBLUP when varying *α*, blending and tuning hyper-parameters for five carcass traits. The five carcass traits include carcass weight (cwt), eye muscle area (ema), adjusted 12 months weight (adj-w12), marbling score (ms) and back fat thickness (bft). The pedigree includes 84,020 animals in total, among which around 20,490 animals have phenotypic records, and 13,800 animals are genotyped for 42,686 SNPs across the genome. The number of animals with both genotypes and phenotypes is 9,072 (Table 2) that are randomly divided into 5 groups (5-fold cross validation). Each set of the five groups is selected as the target samples, and all the phenotyped animals except the target samples were used as the discovery dataset. This five-fold cross validation was used to validate the performance of HBLUP.

## Discussion

HBLUP or ssGBLUP has been widely used in livestock breeding programs [5,6]. The HBLUP method (e.g., BLUPf90) requires hyper-parameters to integrate the information of genomic and pedigree relationship matrices, which should be optimised to maximise the accuracy of genomic prediction [7, 13, 25, 26]. In this study, we evaluated the performance of HBLUP with various hyper-parameters such as blending, tuning and scale factor, using simulated and real Hanwoo cattle datasets.

The scale factor, *α*, can determine the relationship between allele frequency and per-allele effect size. In the simulation, HBLUP accuracy can be the highest when using GRM scaled by the true *α* value used in the phenotypic simulation, indicating that the choice of *α* value is important although this has never been considered as a hyper-parameter in HBLUP. In fact, the performance of HBLUP is shown to vary across the carcass traits in the cattle data used in this study, confirming previous studies reporting that optimal *α* values vary, depending on traits and populations [27–29]. Importantly, using less optimal *α* values may decrease HBLUP accuracy significantly, which should be carefully checked before conducting genetic evaluations, emphasising that the scale factor is not less important, compared to other hyper-parameters such as blending and tuning.

In both simulated and cattle data, blending (*θ* < 1) would not really improve the prediction accuracy except of one cattle trait (EMA, *θ_optimal_* = 0.86). On the contrary, the accuracy would increase more when GRM was blended with higher weights, which is clearly shown in S2 Fig. This is not totally unexpected because richer information can come from GRM (e.g., Mendelian sampling variance within sibs), and blended GRM may lose some of such information. When the mixed model equation is used for HREML or HBLUP [38, 41], a nonpositive definite GRM may cause a numerical problem, for which blending process is essential. This may be one of reasons blending has been an important hyper-parameter in HBLUP. However, the direct AI algorithm can use a non-positive definite GRM without blending (*θ* = 1) and there is a method that can provide positive definite GRM [29]. In any case, we recommend optimising the blending hyper-parameter as the optimal blending can vary, depending on data, in which *θ* = 1 should also be explicitly evaluated.

The tuning process adjusts GRM, accounting for the allele frequencies in the base population, assuming that the founders in the base population are not genotyped but are linked through the pedigree. As expected, the widely use tuning method (tune=1; [26]. implemented in BLUPf90 option 2) could significantly improve the prediction accuracy in the simulated data, indicating that the base allele frequencies are correctly accounted for. However, the improvement caused by tune=1 or 2 was not remarkable in the Hanwoo cattle data. This is probably due the fact that the pedigree information in the real data is not accurate enough to trace the founders, or the genotypes may capture substantial information about the base allele frequencies.

In conclusion, existing hyper-parameters such as blending and tuning in HBLUP are important in general, and their optimal values or options should be properly sought to achieve a reliable genetic evaluation. Depending on data, optimal values can vary, and unnecessary or over-parametrised blending or tuning can produce adverse effects on the prediction accuracy. The scale factor, a novel hyper-parameter to be introduced in HBLUP, should be explicitly optimised to increase the prediction accuracy, given the impact of scale factor is competitive with other hyper-parameters, blending and tuning. We suggest including the scale factor, *α*, in HBLUP as a hyper-parameter.

## Supporting information

Supplementary hyper_parameter

## Acknowledgment

This study is supported by Cooperative Research Program for Agriculture Science and Technology Development (PJ0160992022) from the Rural Development Administration, Republic of Korea.

## Author Contributions

**Conceptualization:** Mehdi Neshat, Soohyun Lee, Buu Truong, Julius H. J. van der Werf, S. Hong Lee.

**Methodology:** Mehdi Neshat, S. Hong Lee.

**Data Curation:** Mehdi Neshat, Soohyun Lee, Buu Truong.

**Formal Analysis:** Mehdi Neshat, Soohyun Lee, Md. Moksedul Momin.

**Investigation:** Mehdi Neshat, S. Hong Lee.

**Software:** Mehdi Neshat, Md. Moksedul Momin, S. Hong Lee.

**Validation:** Mehdi Neshat.

**Visualization:** Mehdi Neshat.

**Resources:** Soohyun Lee, Buu Truong.

**Funding Acquisition:** Soohyun Lee, S. Hong Lee.

**Writing – Original Draft Preparation:** Mehdi Neshat, S. Hong Lee.

**Writing – Review & Editing:** Mehdi Neshat, Md. Moksedul Momin, Buu Truong, Julius H. J. van der Werf, S. Hong Lee.

**Project Administration:** Soohyun Lee, S. Hong Lee.

**Supervision:** S. Hong Lee.

## Supporting information

**S1 Fig. Three steps of the historical population and analysing livestock genomic data simulated using QMSim software.**

**S2 Fig. Adjusting the blending and tuning values of HBLUP (half sib design) for h^2^ = 0.8 with 3000 replications (95% confidence interval).**

**S3 Fig. Adjusting the blending and tuning value for HBLUP with h^2^ = 0.8. with Ne=1000, and 3000 replications (95% confidence interval).**

**S4 Fig. HBLUP accuracy and hyper-parameters.**

**S5 Fig. HBLUP accuracy and hyper-parameters (Tune=1 and Blend=1) using simulated data where** *N_e_* = 100.

**S6 Fig. HBLUP accuracy and hyper-parameters (Tune=1 and Blend=1) using simulated data with** *N_e_* = 1000.

**S7 Fig. The performance of the grid search hyper-parameter adjusting for HBLUP and simulated data using QMSim software**

**S8 Fig. The HBLUP’s prediction accuracy landscape achieved by various tune, blending and alpha values for simulated data (applied the QMSim software) using the grid search hyper-parameter tuning.**

**S1 Table. The best-found hyper-parameters for HREML estimation and HBLUP prediction accuracy for five cattle traits, including adj-w12, BFT, CWT, EMA, and MS.**

## Notes

### Competing Interest Statement

The authors have declared no competing interest.

