## Supplementary hyper_parameter for "A novel hyper-parameter can increase the prediction accuracy in a single-step genetic evaluation"

**SUPPLEMENTARY FIGURES**

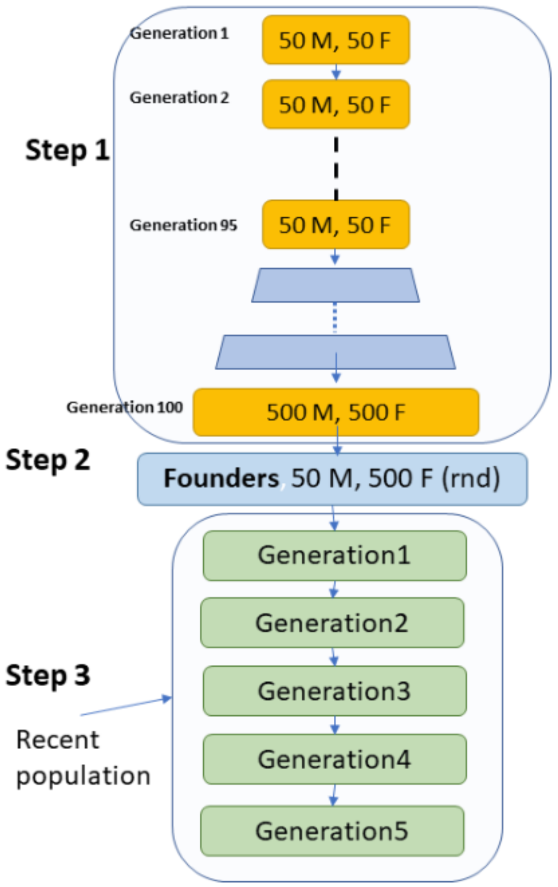


**S1 Fig. Three steps of the historical population and analysing livestock genomic data simulated using QMSim software**. The historical population consists of 100 generations. For the initial 95 generations, the effective population size ($N_e$) keeps fixed at 100 individuals, consisting of 50 females and 50 males. Two offspring are generated using a uniform random of a pair of parents. In the following five generations ($95^{th}$-$100^{th}$), the number of progenies is gradually increased to 1000. Both selection and mating designs are sampled randomly. In the last generation of the historical population (the $100^{th}$ generation, we select randomly 50 males and 500 females recognised as the founders, and each male is mated with ten females and each female produced two offspring (i.e., a half-sib design). The recent population is made in the third step, including five generations with 1000 offspring in each generation.

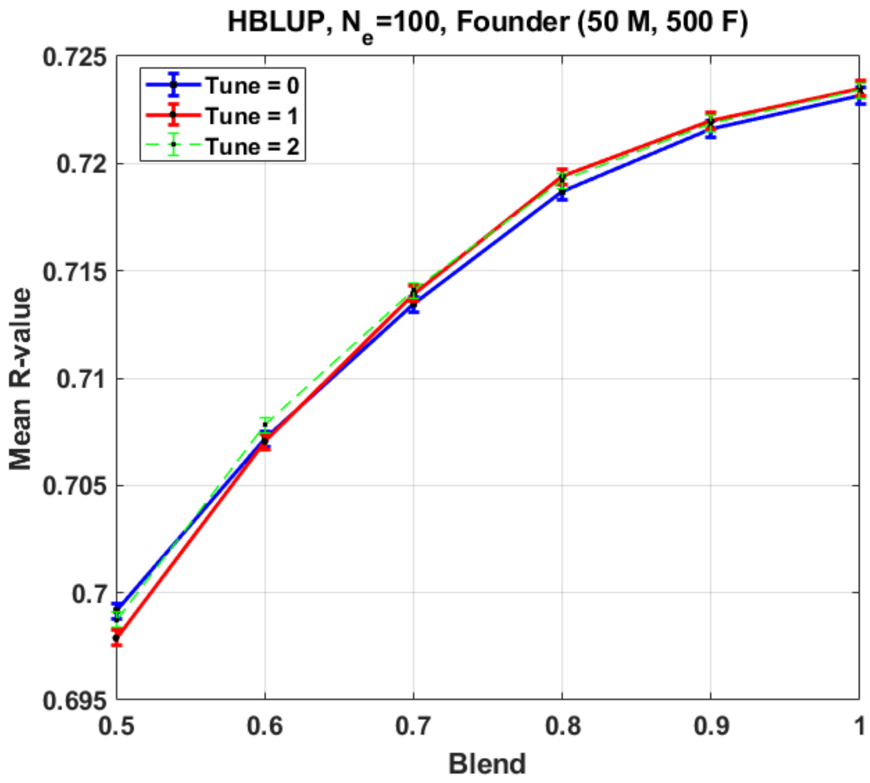


**S2 Fig. Adjusting the blending and tuning values of HBLUP (half sib design) for h2 = 0.8 with 3000 replications (95% confidence interval),** QMSim software is used to simulate the historical population with 100 generations, founders include 50 male and 500 females, the recent population is produced including five generations with 1000 offspring in each generation with effective population size ($N_e$) at 100. Biallelic markers are 9000 and distributed randomly though the genome with the same probability at 0.5 in the first generation. The number of chromosomes is 30. Both selection and mating designs are random. HBLUP: pedigree, genomic information (used in making GRM), target, and discovery sample size are 3550, 2000 (last two generations), 1000 (randomly selected from last two generations), and 4550 respectively.

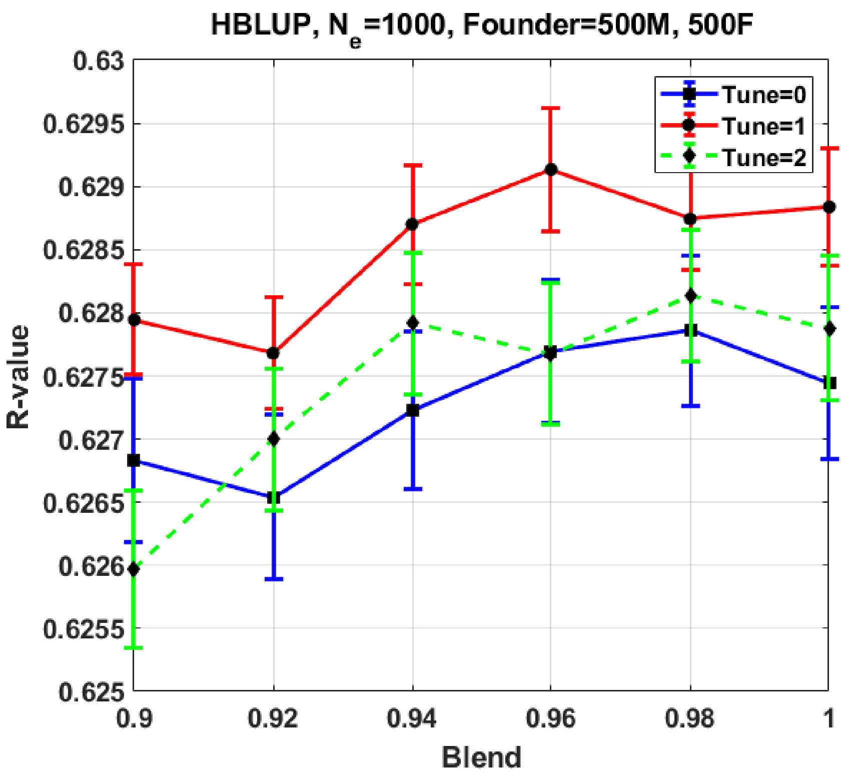


**S3 Fig. Adjusting the blending and tuning value for HBLUP with $h^2$ = 0.8. with Ne=1000, and 3000 replications (95% confidence interval).** QMSim software is used to simulate the historical population with 100 generations, founders include 500 male and 500 females, the recent population is produced including five generations with 1000 offspring in each generation with effective population size (Ne) at 1000, both selection and mating designs are random. HBLUP: pedigree, genomic information (used in making GRM), target, and discovery sample size are 4000, 2000 (last two generations), 1000 (randomly selected from last two generations), and 5000 respectively.

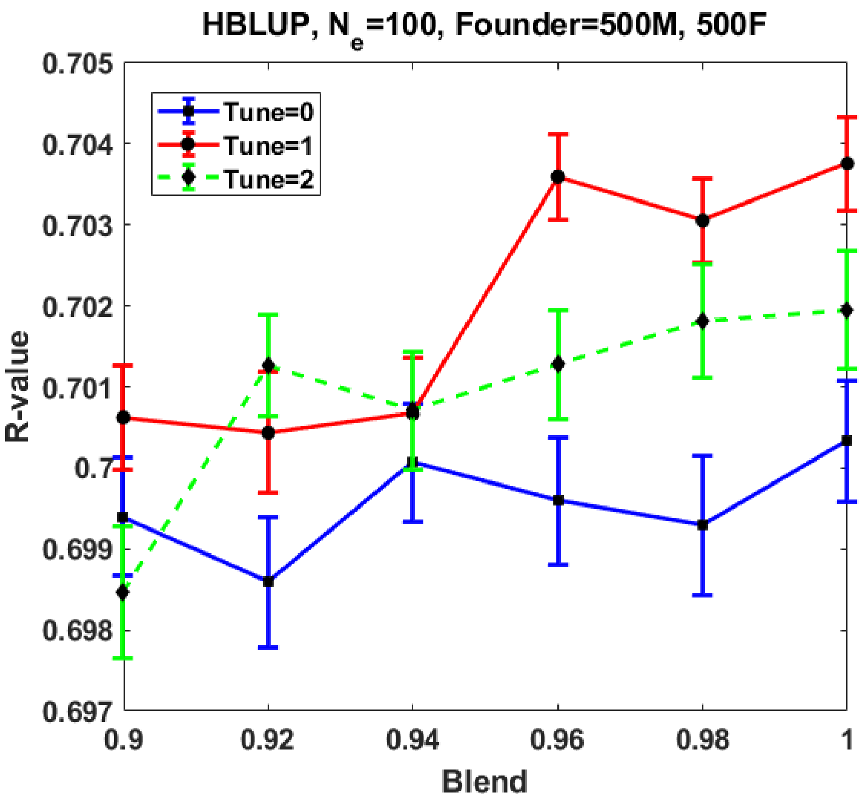


**S4 Fig. HBLUP accuracy and hyper-parameters.**

We simulated genotypes and phenotypes in 3000 replications in which $h^2$ =0.8, $N_e = 100$ for 100 historical generations, (a) a full-sib design (500 male, 500 female), The true $\alpha$ value used in the phenotypic simulation was -0.5. The error bars are 95% CI over the 3000 replications. The pedigree, genotyped individuals (used in making GRM), target, and discovery sample sizes are 4000, 2000 (last two generations), 1000 (randomly selected from last two generations), and 5000 respectively.

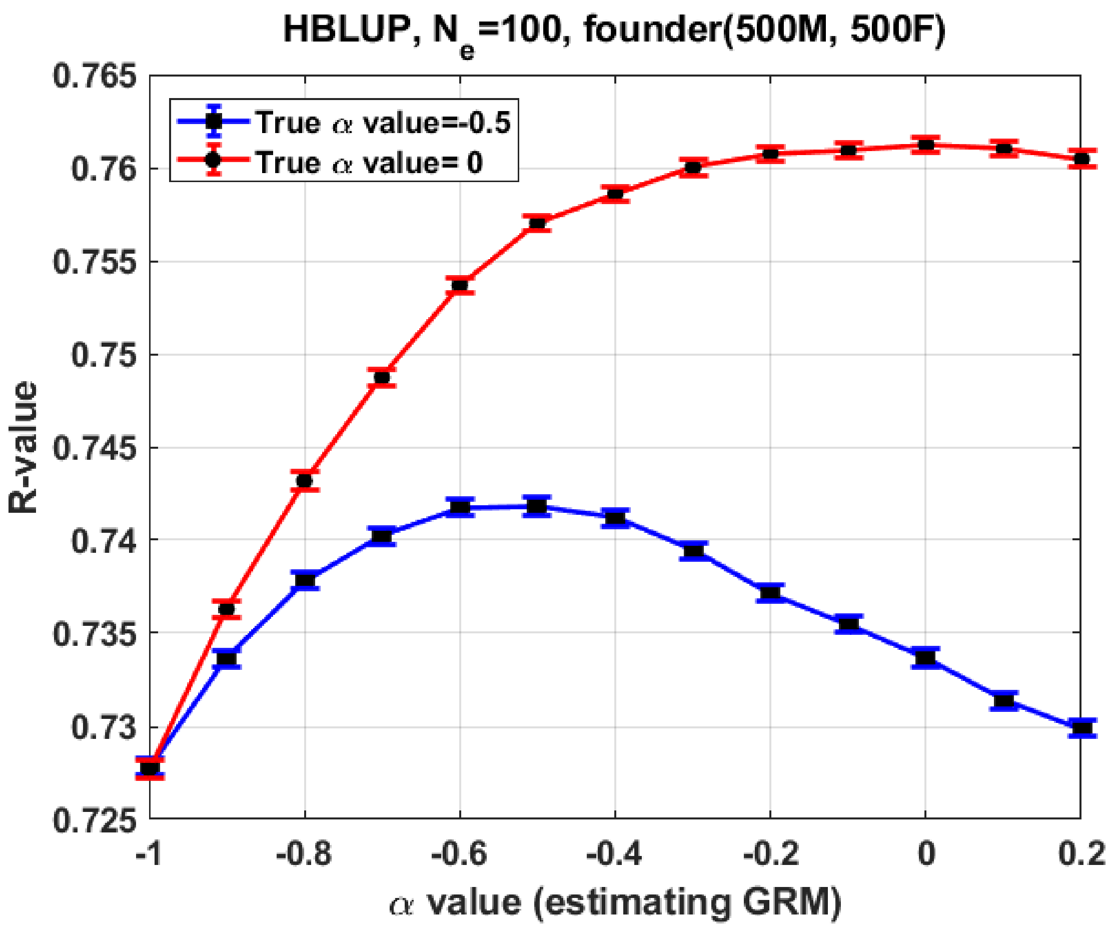


**S5 Fig. HBLUP accuracy and hyper-parameters (Tune=1 and Blend=1) using simulated data where $N_e = 100$.**

Optimal $\alpha$ values can increase the accuracy, indicating that the choice of $\alpha$ is important in HBLUP. We simulated genotypes and phenotypes in 3000 replications in which $h^2$ =0.8, $N_e = 100$ for 100 historical generations, and a full-sib design (500 male, 500 female) used. The true $\alpha$ values used in the phenotypic simulation were -0.5 and zero. The error bars are 95% CI over the 3000 replications. The pedigree, genotyped individuals (used in making GRM), target, and discovery sample sizes are 4000, 2000 (last two generations), 1000 (randomly selected from last two generations), and 5000 respectively.

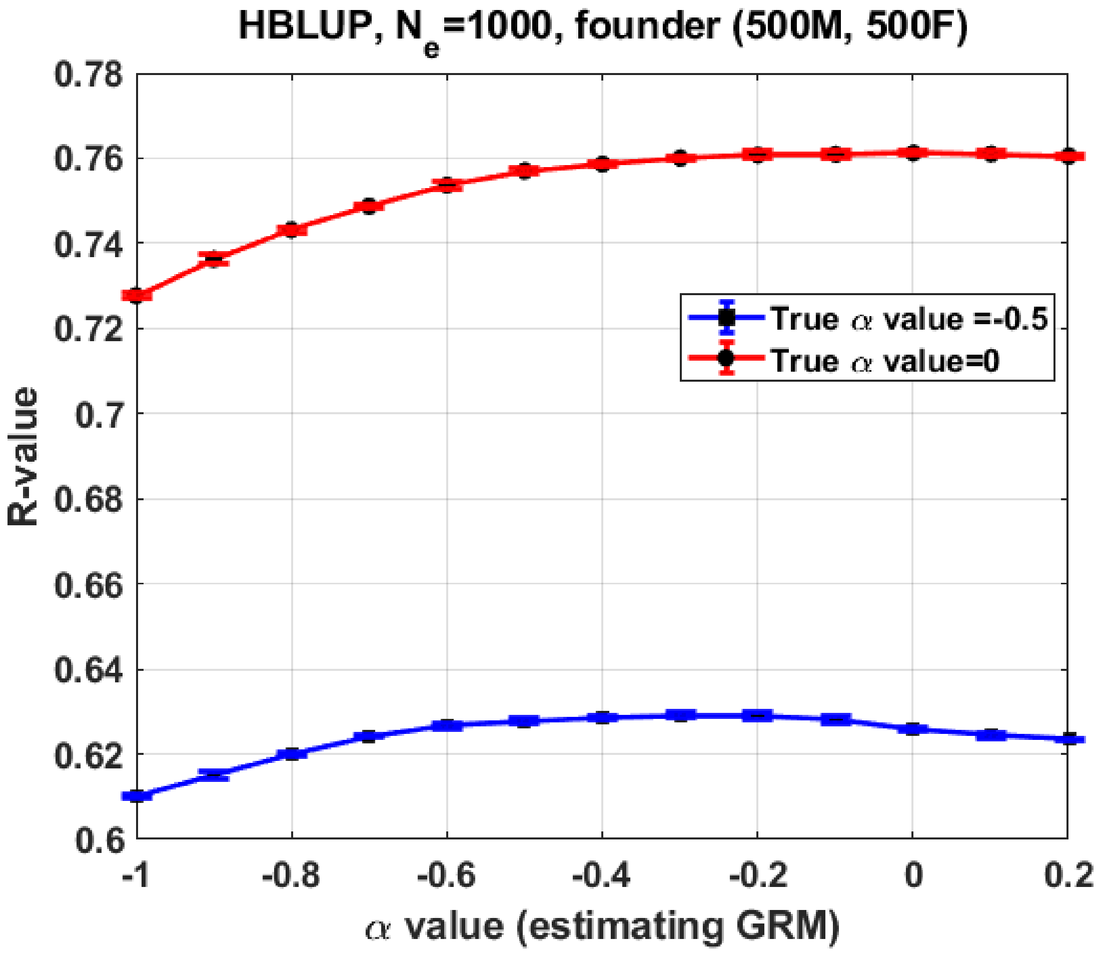


**S6 Fig. HBLUP accuracy and hyper-parameters (Tune=1 and Blend=1) using simulated data with** $N_e = 1000$.

We simulated genotypes and phenotypes in 3000 replications in which $h^2$ =0.8, $N_e = 1000$ for 100 historical generations, and a full-sib design (500 male, 500 female) used. The true $\alpha$ values used in the phenotypic simulation were -0.5 and zero. The error bars are 95% CI over the 3000 replications. The pedigree, genotyped individuals (used in making GRM), target, and discovery sample sizes are 4000, 2000 (last two generations), 1000 (randomly selected from last two generations), and 5000 respectively.

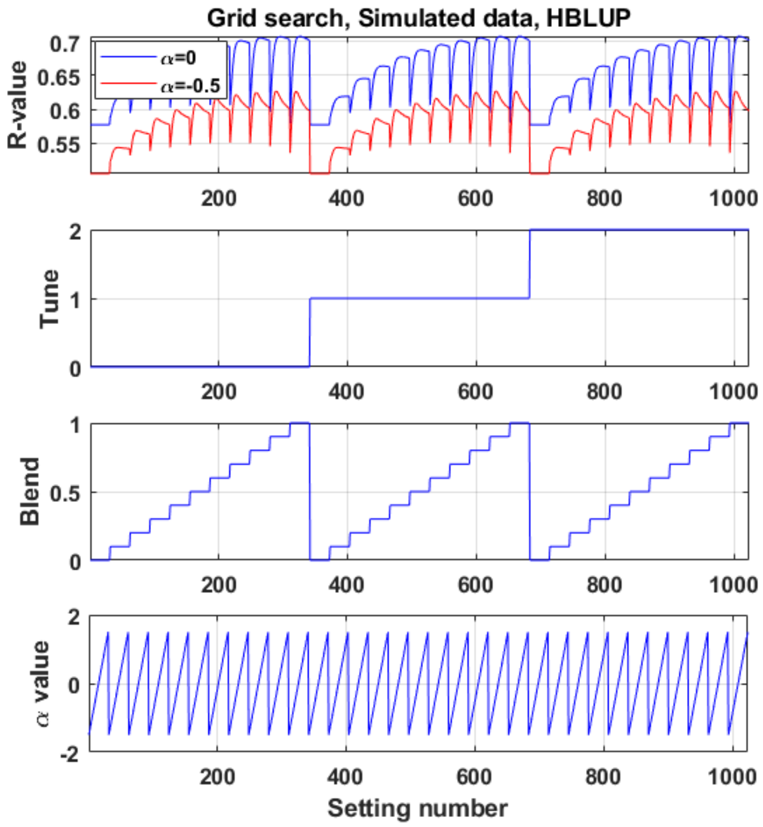


**S7 Fig. The performance of the grid search hyper-parameter adjusting for HBLUP and simulated data using QMSim software.** The value of the hyper-parameters is including tune= (0,1), blend = [0,1], alpha = [-1.5, 1.5], and the applied alpha for simulating phenotypes are -0.5, and zero. The highest HBLUP's prediction accuracy observed for blending=1 and the estimated alpha value is close to the true alpha.

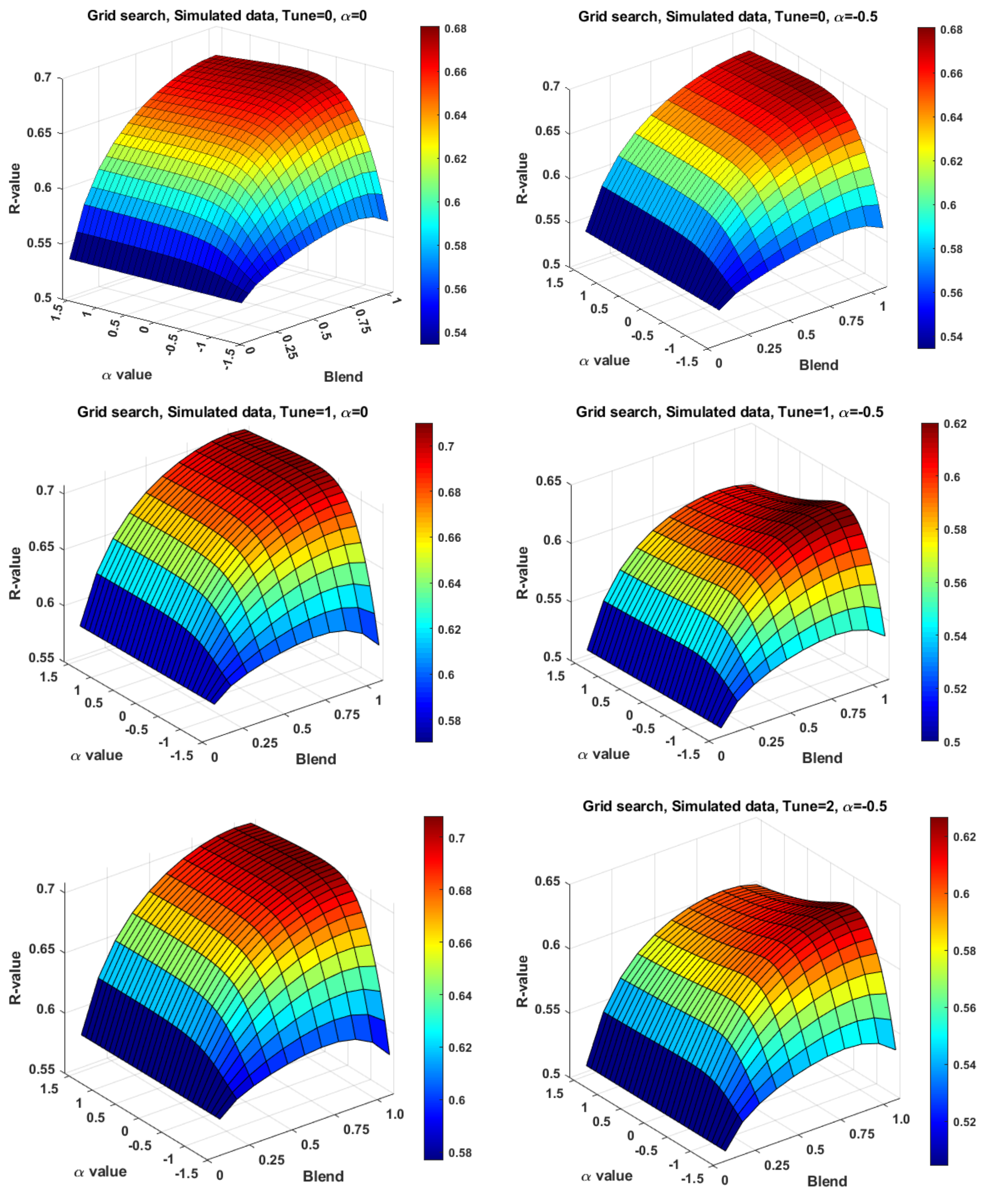


**S8 Fig. The HBLUP's prediction accuracy landscape achieved by various tune, blending and alpha values for simulated data (applied the QMSim software) using the grid search hyper-parameter tuning.** The details of the simulating genotypes are $h^2$ =0.8, $N_e$ = 100, founder (50 male, 500 female (half-sib)), $\alpha$ = -0.5 (left side figures), and zero (right side figures) to simulate the phenotypes, $N_{SNPs}$ = 9000, Chromosome number = 30, target samples = 400, discovery sample size=5150, pedigree sample size=3550. The 5-fold cross validation is used to validate the performance of HBLUP. The dark red areas show the highest prediction accuracy of the HBLUP. R-value is the Pearson correlation coefficient, and it shows the strength of a linear relationship among estimated phenotypes and real values.

**S1 Table. The best-found hyper-parameters for HREML estimation and HBLUP prediction accuracy for five cattle traits, including adj-w12, BFT, CWT, EMA, and MS.**

| **HREML** | | | | | |
|---|---|---|---|---|---|
| | adj_w12 | BFT | CWT | EMA | MS |
| Tune | 2 | 0 | 2 | 2 | 1 |
| Blend | 1 | 1 | 1 | 0.86 | 1 |
| alpha | 0.25 | -0.25 | 0 | 0.5 | 0 |
| **HBLUP** | | | | | |
| | adj_w12 | BFT | CWT | EMA | MS |
| Tune | 0 | 0 | 2 | 1 | 0 |
| Blend | 1 | 1 | 1 | 0.86 | 1 |
| alpha | 0.25 | -0.25 | 0 | 2 | -0.25 |
